# Cortex-specific inversion of visual responses during sleep

**DOI:** 10.64898/2026.01.30.702868

**Authors:** Nicholas G. Cicero, Michaela Klimova, Louis Vinke, Sam Ling, Laura D. Lewis

**Affiliations:** Graduate Program for Neuroscience, Boston University; Boston, USA; Department of Biomedical Engineering, Boston University; Boston, USA; Institute for Medical Engineering and Sciences, Massachusetts Institute of Technology; Cambridge, USA; Department of Psychological and Brain Sciences, Boston University; Boston, USA; Department of Psychology, Northeastern University, Boston, USA; Department of Electrical Engineering and Computer Science, Massachusetts Institute of Technology; Cambridge, USA; Department of Psychiatry, Massachusetts General Hospital; Boston, USA; Athinoula A. Martinos Center for Biomedical Imaging, Department of Radiology, Massachusetts General Hospital; Charlestown, USA

## Abstract

During sleep, we functionally disengage from our external environment. Our eyes close, profoundly reducing visual input to the brain. However, some light passes through the eyelid, and luminance changes are perceived even through closed eyes during wakefulness. Although the relay of sensory information is thought to be gated by the thalamus during sleep, sensory information can still reach the cortex. To elucidate how visual inputs are modulated at each stage of thalamic and cortical processing during sleep, we used simultaneous EEG-fMRI while presenting luminance-modulated visual stimuli to sleeping humans. We discovered that responses to light remained intact in the visual thalamus during N1 and N2 sleep. However, stimulus-evoked responses in early visual cortex were profoundly suppressed, exhibiting an inverted pattern in which high-intensity visual stimulation evoked visual cortical deactivation. These findings suggest a cortical mechanism where inhibitory circuits regulate stimulus-driven deactivation in visual cortex, facilitating sensory isolation during early stages of sleep.

## INTRODUCTION

Withdrawal from the external environment is a defining feature of sleep, as the brain curbs sensory information from entering conscious awareness. Of all sensory stimuli, visual inputs are particularly suppressed since we close our eyes while we sleep, blocking most information from reaching the nervous system. However, light still penetrates the eyelid (*1-2*); during wakefulness, we can see changes in luminance through closed eyes, and high-intensity visual stimulation through closed eyes drives robust responses in the early visual system, including the lateral geniculate nucleus (LGN) and primary visual cortex (V1) (*3*). Although bright light can activate the visual thalamus and cortex through closed eyes, it remains unclear how visual inputs that pass through the eyelid are downregulated within the brain while we sleep.

Subcortical gating mechanisms have long been implicated in sensory disconnection, as thalamic inhibition limits the relay of sensory information to the cortex during sleep (*4-8*). However, while thalamic gating can reduce sensory transmission to the cortex, substantial evidence indicates that sensory stimuli presented during non-rapid eye movement (NREM) sleep are not completely blocked within the thalamus, but rather can reach sensory cortices (*4, 8-21*). Tones and speech presented during sleep evoke responses in the auditory cortex (*9-11, 18-19*) but fail to engage higher-order auditory regions (*20-21*). In the visual system, light also evokes responses in the visual cortex during sleep (*12-16, 19*), altogether suggesting that mechanisms at the level of the cortex are required to modulate sensory processing. Despite these observations, the lack of simultaneous measurements across the thalamus and cortex has left unresolved the subcortical and cortical mechanisms that downregulate visual inputs during sleep.

To investigate the impact of sleep on visual responses in the human thalamus and cortex, we recorded simultaneous electroencephalography (EEG) and functional magnetic resonance imaging (fMRI) responses across the whole brain in 15 human subjects, during wakefulness and the early stages of NREM sleep (N1 and N2) (Fig. 1A).

**Fig. 1.**
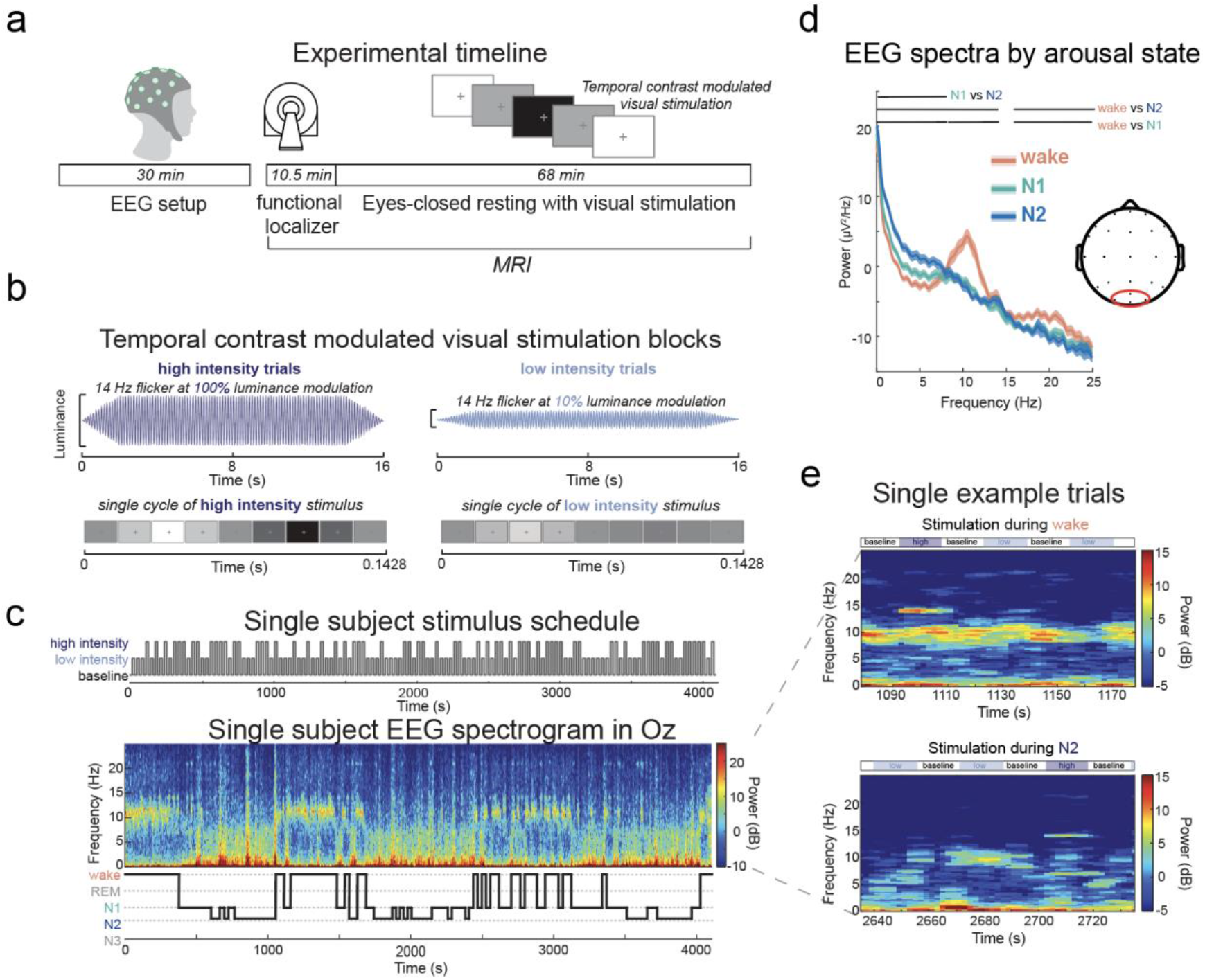
Experimental timeline and example EEG spectrogram during resting scan. **a)** After EEG setup and an anatomical scan, subjects performed three eyes-open runs of a visual functional localizer which included a block-design flickering, full-field checkerboard stimulus, lasting 10.5 minutes total. Subjects then performed an eyes-closed resting scan while a temporal contrast modulated visual stimulus was presented and the subjects performed a self-paced button press task, lasting 68 minutes. Across the resting scan, subjects naturally transitioned in and out of sleep. **b)** Visual stimulation blocks comprised a 14 Hz temporal contrast flicker lasting 16 seconds with either high- or low-intensity luminance modulation. High-intensity trials had 100% luminance modulation, flickering from a completely white to black screen, whereas low-intensity trials had a 10% luminance modulation, flickering between a light and dark gray screen. Luminance modulation of the stimulus linearly ramped up over the first 2 seconds and ramped down over the last 2 seconds of the trial. **c)** An example EEG spectrogram across the entire resting scan from channel Oz, with the corresponding sleep hypnogram from sleep staging (*22*), and pseudorandomized stimulus schedule presented throughout the scan. In this example, the participant transitioned in and out of sleep several times, with awake periods indicated by increased alpha (8-12 Hz) power and sleep periods indicated by increased delta (0.5-4 Hz) power and decreased alpha power. **d)** Mean EEG power spectra (around occipital electrodes, circled in red within inset) across sleep stages. Top lines indicate frequencies with significantly different power between arousal states (*p*<0.05, frequency-binned t-test). Frequency-binned t-tests were corrected for number of frequencies and arousal state pairs tested. **e**) Representative time-frequency representations of scalp EEG (Oz) acquired during wake (top) and N2 sleep (bottom) show single trial responses at 14 Hz. Stimulus schedule for example trials is indicated above each spectrogram, with trial labels within each 16 second long box. Increased 14 Hz power can be seen during high-intensity stimulation in both examples.

## RESULTS

### Stimulus-evoked responses across sleep-wake states in the visual system

We first performed a functional localizer during eyes-open wakefulness, to identify the LGN (Supplemental Fig. 1) and visual cortical regions. We then acquired neural responses as participants rested with their eyes closed and spontaneously transitioned in and out of NREM sleep, while we presented a flickering visual stimulus of different luminance intensities (Fig. 1B). We separated visual stimulation trials by the arousal state in which they occurred (Fig. 1C-E) based on clinical sleep scoring criteria from the EEG data acquired during the scan (Fig. 1C) (*22*).

To determine if visual stimulation evoked different neural responses across arousal states, we mapped visual-evoked fMRI blood-oxygenation-level-dependent (BOLD) responses across visual thalamic and early cortical regions (V1, V2, and V3; Fig. 2A). We first examined responses during eyes-closed wakefulness and found that high-intensity stimulation activated the LGN, V1, and V2, but deactivated V3 (*p*<0.05; Fig. 2B). During wakefulness, LGN and V1 responses were significantly stronger during high-compared to low-intensity stimulation (*p*<0.05), replicating prior work in which visual stimulation through closed eyes elicits robust luminance-dependent responses in the early visual system (*3*). As participants transitioned into N1 and N2 sleep, we observed stimulus-evoked activation within the LGN (*p*<0.05; Fig. 2C-D), similar to the evoked response during wakefulness. Despite responses to light remaining intact within the early visual thalamus, all visual cortical regions displayed strong stimulus-evoked deactivation in N1 and N2 sleep (*p*<0.05; Fig. 2C-D). All visual cortical regions showed a significant decrease in BOLD in N1 and N2 compared to wakefulness (*p*<0.05; Fig. 2C-D). When comparing between N1 and N2, we observed no significant differences across all ROIs responses (all *p*>0.05), indicating similar responses to visual stimulation within the early stages of NREM sleep (Supplemental Fig. 2). Altogether, high-intensity visual stimulation evoked consistent activation in the LGN across arousal states, but strong deactivation of early visual cortex during sleep.

**Fig. 2.**
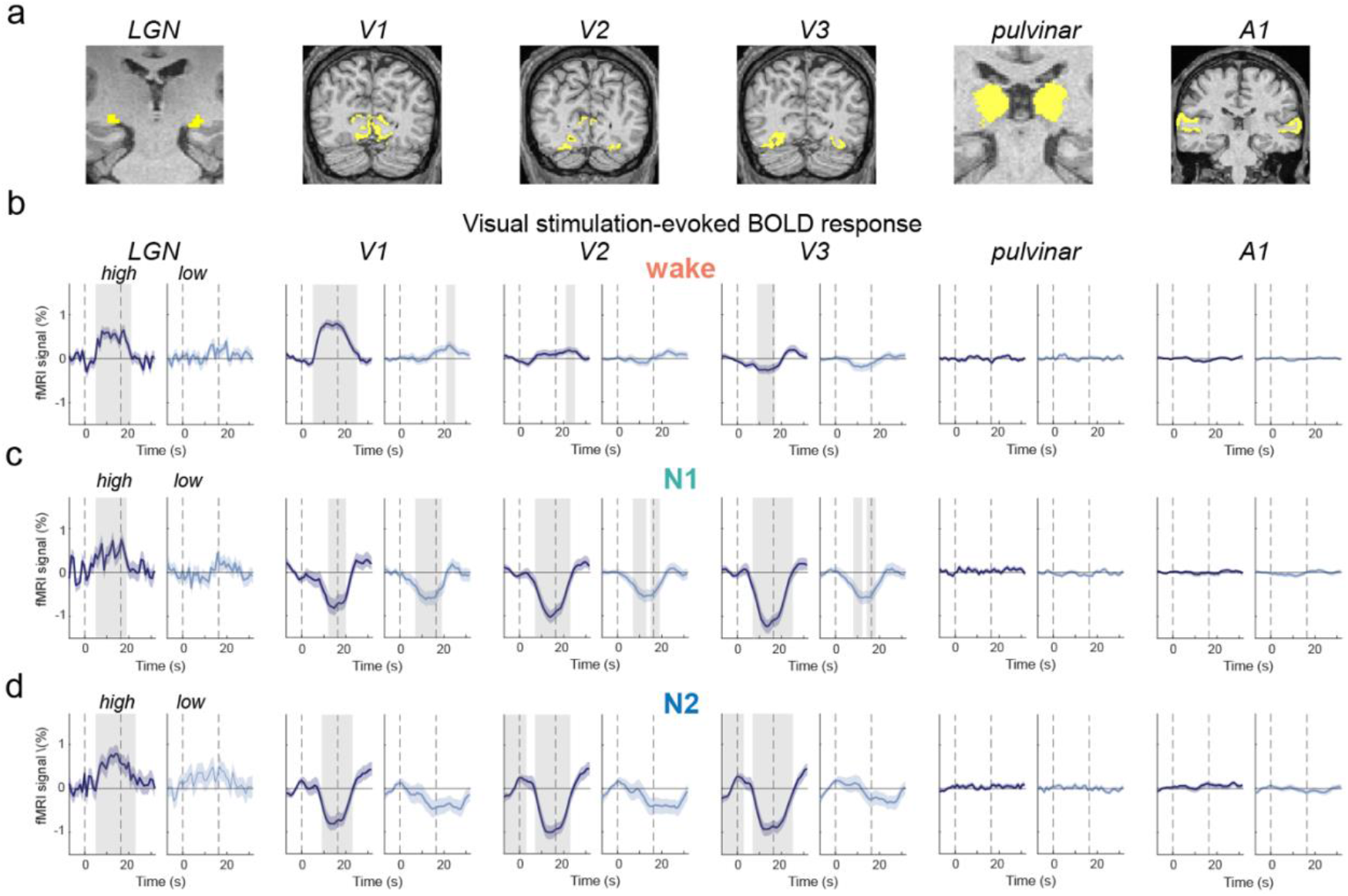
Visual stimulation during sleep produces negative responses throughout early visuocortical regions, despite preserved lower-order visual thalamus activation. **a)** Regions of interest (yellow) are shown in coronal slices in an example subject. **b)** Stimulus-evoked responses by region during wakefulness (high-intensity trials: n=308; low-intensity trials: n=286). During wakefulness, high-intensity stimulation (dark blue) evoked significant BOLD activation in LGN, V1, and V2 (gray shading indicates *p*<0.05), but a significant deactivation in V3 (*p*<0.05). **c)** During N1 (high-intensity trials: n=167; low-intensity trials: n=204) and (**d**) N2 (high-intensity trials: n=145; low-intensity trials: n=137), the LGN displayed significant BOLD activation during high-intensity stimulation, whereas V1, V2, and V3 displayed significant BOLD deactivation to high-intensity stimulation (*p*<0.05). Pulvinar and primary auditory cortex showed no significant changes in activity across all trials and all arousal states during the stimulus presentation (all *p*>0.05). P-values were calculated with sliding window time-binned linear mixed effects models (with subject as a random effect) and corrected for number of time bins, arousal states, and number of ROIs tested using the Benjamini-Hochberg procedure to control the false discovery rate (FDR) (*93*). Grey bars indicate time bins where *p*<0.05. Colored shading is standard error. Dashed vertical lines indicate stimulus onset (left line) and offset (right line). BOLD=blood-oxygenation-level-dependent; LGN=lateral geniculate nucleus; V1=primary visual cortex; V2=secondary visual cortex; V3=tertiary visual cortex; A1=auditory cortex.

We next examined whether this deactivation occurred beyond early visual cortex to investigate the spatial extent of this stimulus-evoked pattern during sleep. Other visual cortical regions along the visual pathway displayed similar deactivation patterns during N1 and N2 sleep (Supplemental Fig. 3). In contrast, the pulvinar nucleus of the thalamus, which receives corticothalamic feedback from higher-order visual regions (*23-24*), showed no stimulus-locked BOLD responses in wakefulness nor NREM sleep (Fig. 2B-D), indicating that stimulation failed to engage higher-order sensory regions during sleep, consistent with prior work (*20-21*). Due to the small size of the LGN, its signal-to-noise ratio is substantially lower than the pulvinar, suggesting that even if undetected stimulus-evoked responses were present in pulvinar, they would be of very small amplitude. We also observed that during wakefulness, V3 and other extrastriate regions already displayed deactivation responses to high-intensity stimuli (Fig. 2B and Supplemental Fig. 3B). As suggested by previous work, eye closure itself may suppress extrastriate cortical responses (*3*); therefore, deactivation in later visual cortical regions during wakefulness likely reflects modulation caused by eye closure. We additionally did not observe any significant change in BOLD within primary auditory cortex (A1), suggesting that stimulus-evoked responses were focal to the visual system (Fig. 2). Taken together, these results demonstrate that during sleep, the progress of visual-evoked activity is strikingly altered: visual stimulation evokes the same positive activity as during wakefulness in early visual thalamus (LGN), whereas stimulation evokes large inverted, negative responses within the visual cortex.

**Fig. 3.**
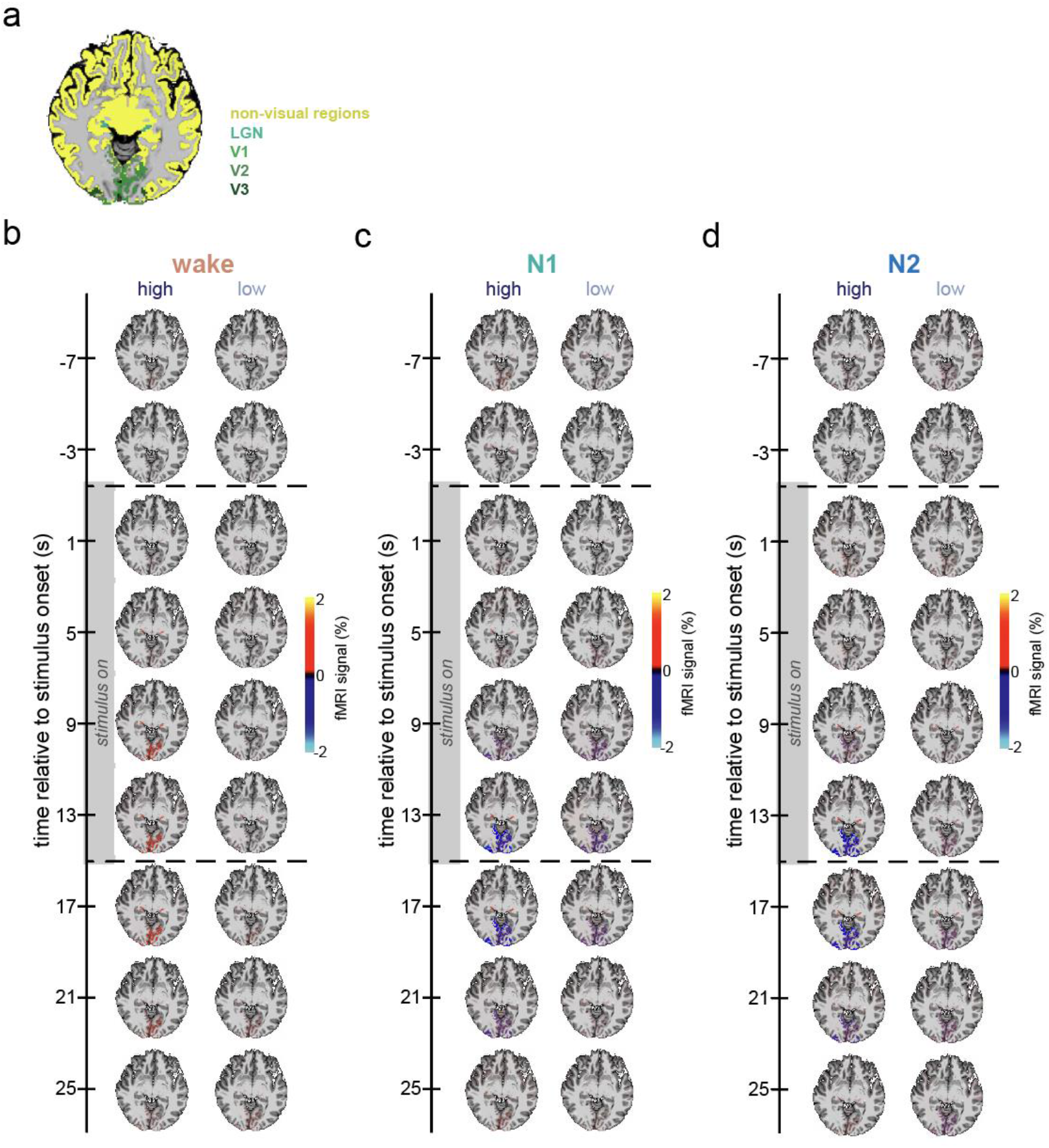
Visual-evoked responses during wakefulness and N1 and N2 sleep are focal to the LGN and visual cortex. Event-triggered average BOLD responses were calculated for each cortical and subcortical region from FreeSurfer’s whole-brain segmentation (*81*). Cortical and subcortical ROI responses were calculated from the average BOLD signal across all voxels within each ROI. **a)** Anatomical ROI masks for visual regions (greens) and all other regions (yellow) are shown outlined in an axial slice. **b-d)** The average ROI responses are displayed within each ROI’s anatomical mask in an example subject’s anatomical space. Whole-brain responses are displayed in four second intervals from −7 seconds to 25 seconds relative to stimulus onset. The displayed responses were thresholded from −2 to 2 BOLD signal change (%). Stimulus presentation is denoted by the grey bar and horizontal dashed grey lines. For trials during wakefulness (**b**), the LGN and V1 more strongly activate during high-compared to low-intensity stimulus presentation. During N1 (**c**) and N2 (**d**) the LGN similarly activates more during high-compared to low-intensity trials while the visual cortex deactivates more during high-compared to low-intensity trials. No other cortical or subcortical regions show a significant BOLD response across arousal states or trial conditions, suggesting the observed response is focal to the visual pathway.

### A global arousal response does not explain visual cortical deactivation

One possible explanation for the observed visual cortical deactivation is that high-intensity stimulation during sleep elicits an arousal response that drives widespread vasoconstriction. Arousals from sleep trigger sympathetic vasoconstriction through interactions between subcortical arousal systems (*25-30*) and the autonomic nervous system (*31-34*), resulting in a global decrease in BOLD across the cortical gray matter accompanied by activation across thalamic nuclei (*35*). First, we did not observe that EEG- or behaviorally-defined arousals occurred more frequently during visual stimulation, as compared to baseline (EEG arousals: *F*(2,39)=0.281, *p*=0.757; behavioral arousals: *F*(2,36)=0.679, *p*=0.513). In addition, our analysis excluded any trials during N1 and N2 sleep where there was an EEG- or behaviorally-defined arousal (see Methods). The evoked responses were also focal to visual cortex, as when examining BOLD responses across the brain we found no significant visual-evoked responses in other cortical or subcortical regions (Fig. 3), nor did we find significant responses in any thalamic nuclei (Supplemental Fig. 4), except for the LGN (Fig. 2), nor across the entire cortical gray matter (Supplemental Fig. 4). Altogether, these findings indicate that the observed visual cortical deactivation cannot be explained by global vasoconstriction induced by a stimulus-driven arousal response. Rather, BOLD deactivation was focal to the visual cortical system, implicating stimulus-driven visual cortical deactivation as a candidate mechanism for sensory disconnection during sleep.

### Stimulus-evoked K-complexes do not mediate visual cortical deactivation

We next investigated whether K-complexes could be a mediating mechanism for the observed visual cortical deactivation. K-complexes are isolated slow waves during N2 sleep that can be triggered by sensory stimulation (*36*), can produce a reduction in cortical BOLD (*37-38*), and are believed to suppress information processing in cortical networks to protect sleep (*4*). Therefore, one possible explanation of the suppressed visual cortical responses is that during sleep, high-intensity stimuli trigger K-complexes and thereby reduce neural activity and BOLD signals. However, prior work has shown that visual stimulation during sleep does not elicit K-complexes (*15*). To investigate this possibility in our dataset, we identified K-complexes throughout our dataset (*n*=133; Supplemental Fig. 5A). We observed that the vast majority of trials (94%) did not contain a K-complex. The frequency of K-complexes was not significantly different across conditions (*F*(2)=1.89, *p*=0.163), such that K-complexes did not occur more frequently during high-intensity (0.033/min) compared with low-intensity stimulation (0.041/min), nor did they occur more frequently during visual stimulation than baseline periods without any stimulation (0.068/min). Furthermore, K-complexes occurred evenly throughout the 16 second duration of all trials rather than at the onset of visual stimulation (Supplemental Fig. 5B). Finally, we found no significant difference in the visual-locked BOLD responses during trials where a K-complex occurred compared to trials without a K-complex (*p*>0.05; Supplemental Fig. 5C). Overall, we observed that visual stimulation was not significantly associated with K-complexes, suggesting that K-complexes do not mediate stimulus-evoked visual cortical deactivation during sleep.

### Known vascular confounds do not explain visual cortical deactivation

To better understand potential mechanisms underlying the inverted response patterns during sleep, we next examined whether vascular confounds might contribute to the negative BOLD responses reported here. Negative BOLD can originate from several possible underlying mechanisms, including neuronal inhibition (*39-40*), the vascular-steal effect (*41-42*), and large increases in oxygen consumption (*43*). The vascular-steal effect posits that a reduction in cerebral blood volume can result from a large amount of blood being diverted to activated areas, leaving adjacent regions that share blood supply with lower blood volume and a negative BOLD response (*41-42*). We did not observe increased BOLD in regions adjacent to the visual cortex during sleep (Fig. 3), and the LGN, which was the only region with a visual-evoked increase in BOLD during sleep, does not share the same vascular network for blood supply as the visual cortex (*44-45*). Thus, the vascular-steal effect cannot account for the observed deactivation. Next, we considered the possibility that the deactivation could be attributed to oxygen consumption from increased neuronal activity far exceeding the oxygen supply, which would result in a net increase in deoxygenated hemoglobin and a negative BOLD response (*44*). It is known that cortical circuits are less excitable during sleep (*46-57*), and we observed that visual stimulation elicited an attenuated steady-state visual evoked potential in occipital EEG electrodes (Supplemental Fig. 6), consistent with prior work (*15*). Additionally, the cerebral metabolic rate of oxygen decreases during sleep (*56-57*), and cerebral blood flow is reduced in response to visual stimuli during sleep (*12-13*), altogether suggesting decreased neuronal activity and less, not more, oxygen consumption during sleep. Finally, mouse studies have shown that neurovascular coupling is similar, or slightly stronger, during NREM sleep (*58*), suggesting that altered neurovascular coupling is unlikely to account for the large observed negative BOLD response.

## DISCUSSION

We show that visual input induces large negative responses in visual cortex in N1 and N2 sleep, suggesting that visual cortical deactivation during NREM sleep is driven by neuronal inhibition (*39-40*). This interpretation is reinforced by evidence that inhibitory interneuron activity can trigger local vasoconstriction (*59-61*), aligning with the inverted visual cortical responses we observed. Notably, the significant deactivation we saw in response to high-intensity luminance during N2 mirrors findings that inhibitory interneuron activity in visual cortex scales with stimulus properties (*62-66*). The mechanisms of luminance processing and brightness adaptation are also proposed to involve feedback inhibition within the visual cortex, directly implicating inhibitory circuits in the processing and adaptation to luminance-modulated stimuli (*67-68*). These inhibitory circuits are themselves strongly modulated by brain state (*66, 69-71*), and sleep is characterized by both increased neuronal inhibition (*72-73*) and a widespread reduction in cortical excitability (*46-55*). For example, during high arousal states (i.e. locomotion), wake-promoting neuromodulators enhance stimulus-driven visual cortical responses by inhibiting local inhibitory circuits (*66*). Sleep may thus produce an inversion of this effect: wake-promoting neuromodulators are globally reduced during sleep (*25*), enhancing the activity of cortical inhibitory circuits, which would suppress visual cortical responses to incoming stimuli that pass through the eyelid. This framework predicts that visual inputs are substantially blocked at the cortical level, consistent with our observation that visual responses during sleep were negative throughout visual cortical regions and failed to engage the pulvinar, and in line with prior work showing that auditory inputs also fail to reach higher-order sensory areas during sleep (*20-21*). The pulvinar receives projections from other regions like the superior colliculus (*23-24*), so contributions from regions other than higher-order visual cortex cannot be ruled out; however, these did not result in detectable stimulus-evoked activity in pulvinar.

Future studies using invasive recording methods with neuronal cell type specificity are needed to test this proposed framework and disentangle the contributions of specific cortical interneuron populations and neuromodulatory systems in sensory gating. A reduction in visual cortical BOLD indirectly reflects a reduction in stimulus-evoked neural activity, but does not prove that information processing itself was impaired at this level. Additionally, the relatively poor temporal resolution of the fMRI scans performed here preclude analyses determining the timing of evoked responses across visual thalamic and cortical regions. Invasive recording studies would be ideally suited to resolve to what extent visual stimuli are encoded and processed during sleep, as well as the relative timing of visual thalamic and cortical responses during sleep. In addition, while we examined multiple thalamic nuclei in this study, the thalamic reticular nucleus (TRN) could not be measured due to its small size (<1 mm) (*74*). Given the proposed role of TRN in sensory gating (*4-8*), one possibility is that TRN contributes to this process through thalamic-mediated modulation of cortical state. Future animal studies could record within the TRN and visual cortex during NREM to clarify its contribution to sensory disconnection mechanisms across the brain.

Further exploration of different stimulus parameters could also aid in revealing the mechanisms underlying this visual cortical deactivation. For example, the relationship between visual-evoked responses from occipital EEG and arousal state depends on the frequency of visual stimulation, such that faster frequencies (>8 Hz) elicit stronger SSVEPs during wake, similar to our observations (Supplemental Fig. 6), while slower frequencies (<5 Hz) instead elicit stronger SSVEP’s during sleep (*15*) and enhance slow wave activity (*16*). These findings suggest that stimulation frequency interacts with the intrinsic oscillatory dynamics of the cortex, enhancing or suppressing the stimulus response depending on arousal state. Inhibitory circuits within visual cortex differentially affect cortical excitation at different stimulation frequencies (*77*); therefore, inhibitory circuits may mediate the interaction between arousal state and sensory stimulation frequency (*15-16*). Along with stimulus frequency, exploring several luminance intensities would more precisely define luminance response functions across arousal states and identify the intensity threshold at which a visual stimulus evokes cortical deactivation during sleep. Future work is also needed to investigate how different individual factors, such as eyelid thickness and skin color, contribute to the extent that sensory information can pervade the eyelid and brain.

Finally, this study focused on changes to visual processing during the early and mid-stages of sleep (N1 and N2), which make up most of the time spent in NREM. This pattern could potentially change substantially in deeper sleep, N3, which is characterized by prominent slow oscillations and increased cortical bistability, in which neurons alternate between depolarized up-states and hyperpolarized down-states. This bistability is believed to suppress sensory and higher-order cortical processing (*78-80*). One proposed manifestation of cortical bistability is the K-complex, an isolated slow wave event that can be trigged by sensory stimulation during N2 sleep (*36*). However, we found that visual stimulation did not evoke K-complexes, consistent with the only prior visual sleep study focused on this question that we are aware of (*15*). Importantly, the absence of stimulus-evoked K-complexes does not imply that the cortex lacks bistable dynamics; rather, it indicates that visual stimulation does not trigger this specific K-complex event, and that the observed visual cortical BOLD deactivation cannot be attributed to stimulus-locked bistable down-states. Although isolated slow wave events (K-complexes) could not explain the stimulus-evoked visual cortical deactivation during N2 sleep (Supplemental Fig. 5), this mechanism could potentially still contribute to sensory suppression when it does occur. In addition, how these sensory-evoked patterns are modulated in deeper sleep remains an open question. It is possible that visual stimulation presented amidst ongoing large-amplitude slow oscillations might engage distinct mechanisms than stimulation presented in lighter sleep. However, our findings suggest that cortical deactivation in response to sensory input emerges in the very lightest stages of sleep, preceding and not necessarily requiring the canonical bistable slow waves of deep sleep.

In conclusion, we find that light input to the sleeping brain causes early thalamic responses that resemble wakefulness, but strong deactivation of early visual cortex. These results implicate cortical inhibition as a key mechanism by which the brain becomes isolated from our sensory environment while we sleep.

## METHODS

### Participants

Data was acquired from a total of 15 healthy participants (9 females, 6 males). Participants were aged 21-43 years (mean=26.5 years), reported normal or corrected-to-normal visual acuity, and were recruited from Boston University and the surrounding community. The day of the study session participants were instructed to restrict caffeine intake before their study participation. All participants provided written informed consent before study enrollment and completed a metal screening form indicating that they had no MRI contraindications. Exclusionary criteria included all MRI contraindications, as well as any sleep, neurological, or psychiatric disorders, or use of sleep-related medications. Participants were reimbursed for their study participation. All aspects of the study were approved by Boston University’s Institutional Review Board.

### Sleep questionnaire

Participants followed their normal sleep schedule prior to the study visit. Participants completed a sleep questionnaire to confirm that no medication that might affect sleep were taken within 24 hours of the study visit. Participants reported their bedtime the night prior, what time they woke up the day of their study, if they took a nap that day, and if they consumed any caffeine the day of their study. The average bedtime the night prior to the study was 9:53 PM (range: 9:00 PM – 3:00 AM) and the average time they woke up was 7:14 AM (range: 4:00 AM – 10:00 AM). No subjects reported napping the day of their session.

### Experimental design

Participants completed a simultaneous EEG-fMRI scan that lasted approximately two hours. All sessions began between 1:00PM to 2:30PM, to serve as an afternoon nap opportunity. Sessions began with an anatomical scan, followed by three consecutive runs of a functional localizer with their eyes open, each run lasting 208 seconds (∼3.5 minutes) (Fig. 1A). The visual stimulus for the functional localizer contained a full field flickering grating stimulus (flicker frequency = 10 Hz; diameter = 6.0°) lasting 16 seconds with a centered circle (diameter = 0.8°). Within the centered circle, letters rapidly appeared one at a time with a new letter appearing every 200 ms. Participants were instructed to press a button whenever the letters ‘J’ and ‘K’ appeared within the centered circle. During the localizer blocks, the display alternated between a full field flickering grating stimulus (16 second duration) and a full field non-flickering display at median luminance value (16 second duration). Participants completed 12 total blocks (6 flickering, 6 non-flickering) with an extra non-flickering block at the beginning of the run. At the end of each functional localizer run, participants were asked to report their wakefulness level (1: drowsy/asleep, 2: somewhat drowsy, 3: mostly awake, 4: fully awake). Across the entire dataset, 61% of participants reported being mostly awake and 39% reported being fully awake, with no participants reporting drowsiness or sleep. Along with reporting their wakefulness level, real-time eye monitoring was carried out using an EyeLink1000 to ensure subjects were awake and kept their eyes open for the duration of the functional localizer scans.

Participants then completed a long scan with a sleep opportunity while a block-design temporal contrast modulated visual stimulus was presented. High- and low-intensity trials were pseudo-randomly ordered, with all high- and low-intensity trials interleaved with a baseline event. All runs consisted of the same number of high- and low-intensity trials, but the order was randomly shuffled with baseline trials interspersed in between all stimulation trials. Additionally, the presentation of stimuli was independent of sleep-wake stage. The visual stimulus began and ended with a non-flickering baseline trial. Each run contained 128 trials (64 high-intensity, 64 low-intensity) interspersed with 128 baseline events, lasting a total of 4096 seconds (∼1 hour and 9 minutes). Throughout the scan, participants were instructed to press a button after each full breath cycle (1 inhale, 1 exhale). This button task was chosen as a behavioral measure of responsivity that has been previously used to track behavioral arousal state during rest (*35, 81*). Data from the button task was unavailable for one subject due to hardware malfunction. For each run, participants were instructed to keep their eyes closed the entire time and were told that they were allowed to fall asleep. To ensure participants kept their eyes closed for the duration of the resting-state scan, real-time eye monitoring was carried out using an EyeLink1000. Experimenters manually scored videos for eye openings longer than one second. No eye openings were observed across all resting-state scans; therefore, no trials were excluded for eye openings.

### Apparatus & stimuli

Stimuli were generated using custom software written in MATLAB (version 2019b) in conjunction with Psychtoolbox (*82*). Participants viewed stimuli that was back-projected onto a screen set within the MRI scanner, using a ProPIXX DLP LED (VPixx Technologies) projector system (refresh rate: 60 Hz; minimum luminance: 1.2 cd/m2; maximum luminance: 2507.9 cd/m^2^). Photometer measurements (model LS-100; Konica Minolta) carried out before the study were used to verify the linearity of the display (1 digital-to-analog conversion (DAC) step = 9.835 cd/m^2^). These measurements were used to calculate the stimulus luminance and were acquired from the inner-facing side of the back-projection screen while positioned within the MRI scanner bore. This was done to best account for the attenuation in luminance due to back-projection screen characteristics.

During the resting-state run, participants were instructed to keep their eyes closed while shown a full screen flickering display (17 degrees of visual angle) with no spatial contrast (Fig. 1A). The full field flicker was presented in a block design with three trial types (baseline, high, and low temporal luminance contrast), with each trial lasting 16 seconds. In the *baseline* trials, the full field display was a constant median luminance with no luminance modulation. During *high-intensity* trials, the full field display flickered with an amplitude envelope of 100% around the middle luminance value (black to white). For *low-intensity* trials the full field display flickered with an amplitude envelope of 10% around the median luminance value (light gray to dark gray). For high- and low-intensity trials the full luminance-modulated cycle had a frequency of 7 Hz; however, reversals from black to white or white to black occurred twice per cycle, resulting in an effective stimulus frequency of 14 Hz (Fig 1A). Additionally, to gradually start and stop the visual stimulation, the amplitude envelope of luminance modulation ramped linearly over the first two seconds of the stimulus and then ramped down over the last two seconds. This stimulus is identical to our previous work (*3*), but the task duration was extended to last for the entire resting-state scan.

### MRI data acquisition

All neuroimaging data were acquired using a research-dedicated Siemens Prisma 3T scanner using a Siemens 64-channel head coil. A whole brain anatomical scan was acquired using a T1-weighted multi-echo MPRAGE (1 mm isotropic voxels; field of view (FOV) = 192 x 192 x 134 mm, flip angle (FA) = 7.00°, repetition time (TR) = 2200 ms, echo time (TE) = 1.57 ms). Functional scans were acquired using T2*-weighted gradient echo planar imaging (2 mm isotropic voxels; FOV = 104 x 104 x 70 mm, FA = 64.00°, TR = 1000 ms, TE = 30 ms, SMS factor = 5, GRAPPA acceleration = 2, slices = 70).

### MRI preprocessing

T1-weighted anatomical data were analyzed using the standard “recon-all” pipeline provided by the FreeSurfer neuroimaging analysis package (*83*), generating cortical surface models, whole brain segmentations, and cortical parcellations. Visuocortical regions of interest (ROI) were extracted from a Bayesian mapping approach (*84*).

Functional BOLD time-series data were slice-time corrected, motion corrected, and then registered using boundary-based registration between functional and anatomical spaces (*85*). To optimize spatial precision of experimental data, no volumetric spatial smoothing was performed (full-width half-maximum 0 mm). To achieve precise alignment of experimental data within the session, cross-run within-modality robust rigid registration was performed, using the middle time point of each run (*86*). BOLD time-series data were high-pass filtered with a cutoff of 0.01 Hz and converted to units of percent signal change.

### Regions of interest (ROIs)

Visuocortical labels were initially segmented in subject anatomical space from a Bayesian mapping approach (*84*). To obtain LGN labels and visuocortical labels, a general linear model (GLM) was computed from the three runs of the functional localizer. Each subject’s statistical output from the GLM was registered to the subject’s anatomical image using the boundary-based registration transformation (*85*) between the functional and anatomical spaces and voxels within the LGN were manually labeled from the t-values from the GLM output. An example subject’s LGN labels with the corresponding GLM activation map is provided in Figure S1. Visuocortical voxels from the initial anatomical segmentation (*84*) were thresholded such that voxels had to appear in the functional localizer to be included in the final V1/V2/V3 ROI. The pulvinar and other thalamic nuclei were segmented in subject’s individual anatomical space using a Bayesian segmentation approach (*87*) to examine thalamic nuclei other than the LGN. All other cortical and subcortical ROIs from the whole-brain segmentation were extracted in subject’s individual anatomical space from the standard “recon-all” pipeline provided by the FreeSurfer (*83*). These thalamic nuclei were anatomically segmented, whereas LGN was functionally segmented; this is both because functional localizers do not exist for most thalamic nuclei, and also because converging evidence indicates high accuracy of segmentation of most larger thalamic nuclei (*75-76*), whereas the smaller and more distant LGN benefits from functional localization. Additionally, we did not include ROIs along the dorsal visual stream (e.g. MT, MST) because reliable localization of these regions is achieved through functional localizers with dedicated task designs, which we did not collect due to time constraints. All ROIs were then transformed from subject anatomical to functional space using the robust rigid transformation matrix with trilinear interpolation and thresholded such that functional voxels that were at least 70% filled by the original anatomical mask were included.

### EEG acquisition and preprocessing

EEG was collected using a 32-channel MR-compatible EEG cap and BrainAmp MR amplifiers (Brain Products GmbH, Gilching, Germany). To ensure safety, subjects were not allowed to proceed to EEG-fMRI recordings unless the impedance of all channels was below 100 kOhms. This impedance cutoff was for participant safety, not for data quality. EEG was collected at a sampling rate of 5000 Hz, synchronized to the scanner clock, and referenced to channel FCz.

EEG preprocessing began with gradient artifact correction where gradient artifacts were removed through average artifact subtraction (*88*) over the average of the 16 TRs comprising each trial. We performed gradient artifact correction across windows of data marked by the beginning and end of each trial to avoid trials with and without visual stimulation influencing the average template that was subtracted from the epoch. Electrodes were then re-referenced to the common average of EEG channels, excluding channels on the face and cheeks. The EEG data was then downsampled to 500 Hz. Carbon-wire loop regression was then performed to remove the ballistocardiogram artifact (*89-90*).

### Sleep scoring

Manual sleep scoring was performed according to the established guidelines of the American Academy of Sleep Science (AASM; *22*). EEG data from F3/F4, C3/C4, and O1/O2 and EOG from E1/E2 were visualized in 30 s epochs together with EMG. Following AASM guidelines, epochs were scored as wake if greater than 50% of the epoch contained 8-12 Hz alpha rhythm in occipital electrodes, as well as irregular eye movements associated with high EMG muscle tone. Epochs were scored as N1 if the occipital alpha rhythm was attenuated and replaced by low amplitude, mixed frequency activity for greater than 50% of the epoch, as well as slow eye movements. Finally, epochs were scored as N2 if one or more K-complexes unassociated with arousals or sleep spindles were observed during the epoch. Sleep scoring revealed that on average participants spent 45% of the time awake (range = 2% to 80%; std = 23%), 32% in N1 (range = 8% to 53%; std = 13%) and 21% in N2 (range = 0% to 64%; std = 21%). Wake epochs were on average 150 seconds long (range = 30s to 1950s), N1 epochs 86 seconds long (range = 30s to 631s), and N2 epochs 149 seconds long (range = 30 to 961s). On the button press task we observed that trials scored as wake had an average click rate of 15.98 clicks/min (std = 8.61 clicks/min), while trials during N1 had an average click rate of 7.82 clicks/min (std = 5.38) and those during N2 had an average click rate of 0.86 clicks/min (std=2.84 clicks/min). Since our resting-state run was a little over an hour and was performed in the daytime, we did not expect subjects to enter N3 or REM. A breakdown of the sleep scoring for each subject is displayed in Table S1.

### Sleep micro-architecture and arousals

To investigate sleep micro-architecture dynamics, we identified K-complexes and arousals from the EEG data. Sleep arousals were visually inspected and manually labeled following the AASM guidelines (*22*). We observed 267 EEG-defined arousals throughout the dataset (baseline trials = 140; low-intensity trials = 55; high-intensity trials = 72). K-complexes present in frontal or central channels were visually inspected and manually labeled using Visbrain (*91*) according to the AASM definition of K-complexes (*22*). The onset of K-complexes was defined as 550 ms prior to the negative peak since this is typically the time for the sharp negative wave (*92*) and the negative peak time was identified using MATLAB’s *findpeaks* function. We observed 133 K-complexes throughout the dataset baseline trials = 65; low-intensity trials = 38; high-intensity trials = 30). We computed a linear mixed-effects model to test if K-complexes occurred more frequently during specific trials, with a random effect of subject.

As an additional metric to detect changes in arousal state evoked by the visual stimulation, we identified behavioral arousals from the button press task. Based on a previous study that used the same behavioral task, we defined a behavioral arousal as a button press after at least 20 seconds of unresponsiveness (*35*). We observed 242 behavioral arousals throughout the dataset (baseline trials = 116; low-intensity trials = 56; high-intensity trials = 70).

### Analysis of SSVEP and BOLD across clinically scored arousal states

To assess the impact of arousal state on visual-evoked responses in the EEG and BOLD data, each trial of the block-design visual stimulus was categorized according to the sleep stage that it occurred in. Any trials where an EEG-or behaviorally-define arousal occurred, or sleep scoring changed from wake-to-NREM sleep or NREM sleep-to-wake during the trial, were discarded to ensure that the entire trial was within a consistent arousal state. In total, 363 trials were excluded. Transitions from wake to N1 constituted 133 of these excluded trials, and transitions from N1 to N2 constituted 60 if these trials. The other excluded trials constituted transitions from either N1 or N2 back to wake. A summary of the number of visual stimulation trials within each sleep stage for each subject is provided in Table S1.

To assess the electrophysiological visuocortical response to the visual stimulus across arousal states, the EEG spectrum in occipital electrodes was computed. For each trial, EEG data from four occipital channels (Oz, O1, O2, POz) were averaged together and then segmented into 16 second epochs based on the stimulus schedule and then linearly detrended. Artifact trials (<5% across all participants) were identified as epochs whose variance and absolute maximum amplitude were more than three scaled median absolute deviations (MAD) from the median (using MATLAB’s *isoutlier* function).

The spectrum of the EEG data was computed for each trial (using MATLAB’s *pspectrum* function) and the baseline trial preceding it. To confirm that trials during wakefulness, N1, and N2 reflected distinct arousal states, EEG spectra for high- and low-intensity trials were first averaged within each subject (Fig. 1D). Then significant differences in the EEG spectra across arousal states were compared using paired t-tests at each frequency and all p-values were corrected for the number of frequencies tested using the Benjamini-Hochberg procedure to control the false discovery rate (FDR; *93*).

We calculated the average stimulus-locked EEG multi-taper spectrogram (tapers = [3 5], moving window = [4 0.5]) for all trials separated by arousal state (Fig. S6A). To visualize the topography of the SSVEP, for each electrode we extracted ± 0.5 Hz around the stimulus flicker frequency (14 Hz) and averaged the power across the 16 second stimulus presentation (Fig. S6A). To visualize the average SSVEP in the time domain, for each trial we bandpass filtered the data ±0.5 Hz around the stimulus flicker frequency (14 Hz), baseline normalized each trial to the average amplitude 4 seconds prior to the stimulus onset, and then averaged all trials of the same trial type and arousal state (Fig. S6B). To isolate the steady-state visual evoked potential (SSVEP), EEG spectra on each trial were normalized by the preceding baseline trial where there was no temporal contrast. If the trial and preceding baseline trial were both within the same arousal state, the EEG spectra for the baseline trial was subtracted from the trial EEG spectra to produce “difference” spectra that reflected only the visual stimulus-evoked oscillations, otherwise the trial was excluded. Then “difference” spectra across trials of the same stimulus intensity and within the same arousal state were averaged together. To compare the SSVEP across stimulus intensity and arousal state, we extracted the EEG power from the “difference” spectra ± 0.5 Hz around the stimulus flicker frequency (14 Hz) (Fig. S6C). A linear mixed effects model was computed to assess the main effects of trial type and arousal state and their interaction on 14 Hz SSVEP power, while controlling for the random effect of subject. A 2-way ANOVA was computed from the fixed effects from the linear mixed effects model. Paired t-tests across trial type and arousal state were computed and p-values were corrected for the number of comparisons using Bonferroni correction. To visualize the steady-state visual-evoked potential, EEG data for each trial was bandpass filtered ± 0.5 Hz around the stimulus flicker frequency (14 Hz) and baseline normalized to the average amplitude 4 seconds prior to stimulus onset. Then the bandpass filtered signal was averaged across all trials within the same arousal state to visualize the average steady-state visual-evoked potential.

An event-triggered average of the BOLD data was computed for each stimulus intensity condition (low and high) per ROI. The BOLD time-series for each ROI per run was separated by the high- and low-intensity trials and then separated by arousal state. Trials were only included if a given arousal state was sustained for the entire duration of the stimulus presentation. Additionally, any trials with a framewise displacement greater than 0.5 mm at any point in the epoch were excluded. Trials were not averaged within subject, but rather all trials of a given type were averaged together and baseline normalization was performed by subtracting the average BOLD −8s to −1s relative to visual stimulation onset. To detect if BOLD responses were significantly different from baseline, sliding window time-binned linear mixed effects models were calculated across 5 TR bins with a 1 TR step and p-values were corrected for the number of bins, number of arousal states, and number of ROIs tested using FDR (*93*). Significant differences in BOLD between high- and low-intensity trials were identified by computing a sliding window time-binned linear mixed effects model across 5 TR bins with a 1 TR step and p-values were similarly corrected for the number of bins, number of arousal states, and number of ROIs tested using FDR (*93*). To account for the variable number of trials per arousal state per subject, a random effect of subject was included in all linear mixed effects model. Event-triggered averages were computed for all visuocortical ROIs (Fig. S3), thalamic nuclei (Fig. S4), and all other cortical and subcortical ROIs from the whole-brain segmentation (Fig. 3 & S4). To summarize the BOLD responses across arousal states, we calculated the average BOLD magnitude during the 16 second stimulus presentation for each ROI for all trials. We then averaged each ROI’s BOLD magnitude for each trial type and condition within each subject, before calculating the average and standard error across subjects (Fig. S2). We additionally calculated the difference in BOLD magnitude between high- and low-intensity stimulation per ROI per subject (Fig. S2).

To investigate BOLD changes evoked by a K-complex, BOLD was extracted from three seconds before to 20 seconds after the onset of the K-complex. Any trial with a framewise displacement greater than 0.5 mm at any point in the epoch was excluded, as were any trials associated with an EEG-defined arousal. All events were averaged together, and baseline normalization was performed by subtracting the average BOLD −8s to −1s preceding the K-complex. To detect if K-complex-evoked BOLD responses were significantly different from baseline, sliding window time-binned linear mixed effects models were calculated across 5 TR bins with a 1 TR step and p-values were corrected for the number of bins and number of ROIs using FDR (*93*). To investigate the differences in BOLD responses during NREM sleep with and without the occurrence of a K-complex, we separated trials during N2 sleep in which a K-complex occurred during the 16s stimulations presentation from trials without a K-complex (Fig. S5). Significant differences in BOLD between N2 trials with and without a K-complex were identified by computing a sliding window time-binned linear mixed effects model across 5 TR bins with a 1 TR step and p-values were corrected for the number of bins and number of ROIs tested using FDR (*93*).

## Data availability

The data that supports the findings of this study will be shared publicly upon acceptance.

## Code availability

This study used freely accessible software: FreeSurfer, FSL, AFNI, and EEGLAB. BASH and MATLAB functions were used to call these standard tools for data analysis.

## Acknowledgements

This work involved the use of instrumentation supported by the NSF Major Research Instrumentation grant BCS-1625552. We acknowledge the University of Minnesota Center for Magnetic Resonance Research for use of the multiband-EPI pulse sequences. Data was analyzed on a high-performance computing cluster supported by the ONR grant N00014-17-1-2304. We thank Stephanie Anakwe, Banban Tan, Emilia Schimmelpfennig, Joseph Licata, Shruthi Chakrapani, and Stephanie McMains for assistance with data collection, and members of the SL laboratory and LDL laboratory for helpful feedback on the manuscript. This work was funded by National Institutes of Health [grant number R01EY028163, U19NS128613, F31NS139696], Sloan Fellowship, McKnight Scholar Award, Pew Biomedical Scholar Award.

## Author contributions

Conceptualization: NGC, MK, LV, SL, LDL; Methodology: NGC, MK, SL, LDL; Investigation: NGC; Visualization: NGC, SL, LDL; Funding acquisition: SL, LDL; Project administration: NGC, SL, LDL; Supervision: SL, LDL; Writing – original draft: NGC; Writing – review & editing: NGC, MK, LV, SL, LDL

## Additional information

Supplementary Information is available for this paper.

