## Supplemental Materials for "Cortex-specific inversion of visual responses during sleep"

### Supplementary Information

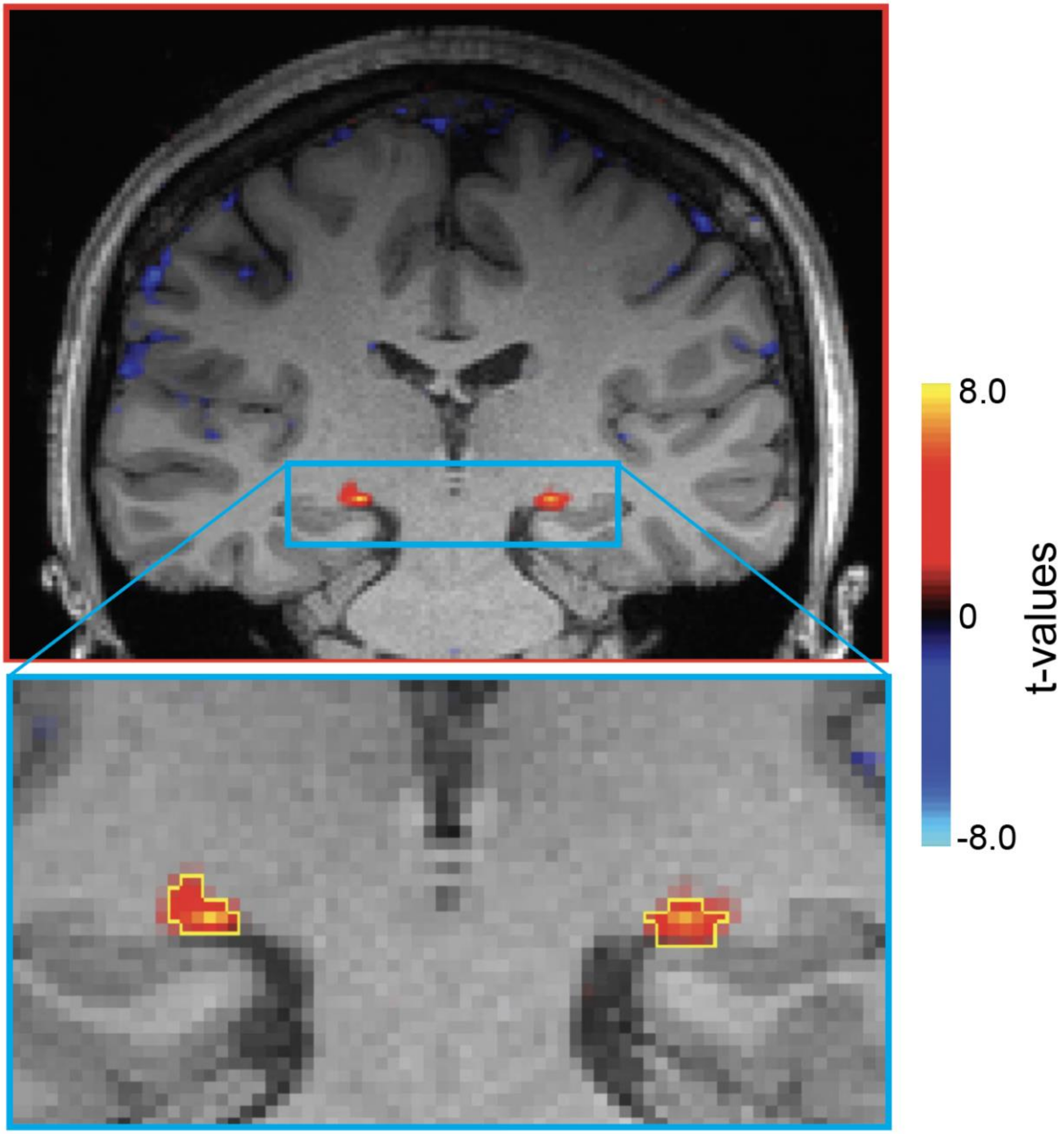


**Fig. S1. LGN ROI drawn from functional localizer.** Example subject's functional localizer GLM results in coronal slice. Top: Whole brain results. Bottom: Zoomed-in functional map with manual LGN ROI voxel selection outlined in yellow. Statistical map thresholded from -8 to 8, excluding t-values between -2 to 2.

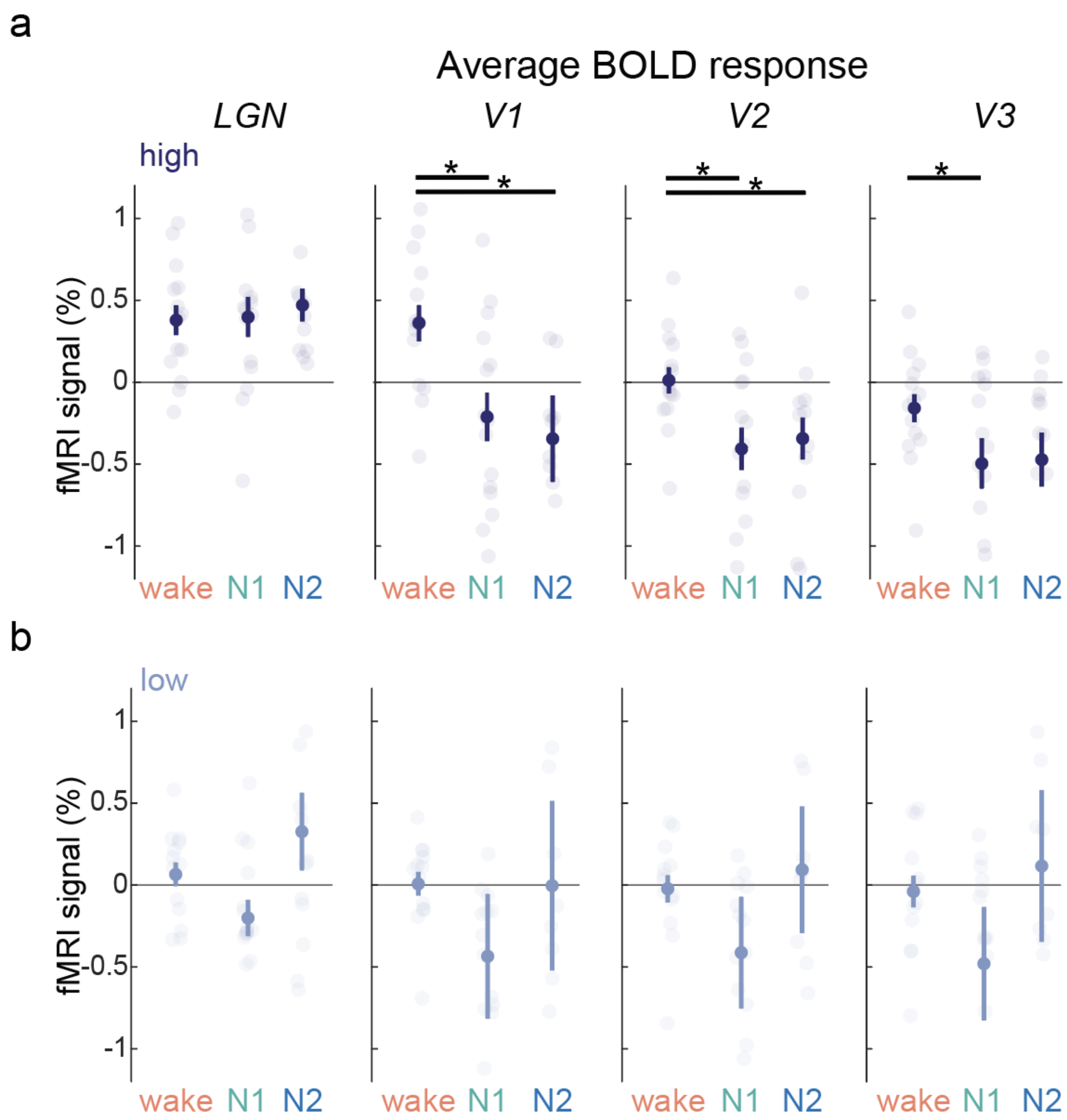


**Fig. S2. Summary of BOLD responses during visual stimulation across sleep-wake states. a-b)** For each ROI, the average BOLD response across stimulus presentation (0-16 seconds) was calculated per trial within each subject. Average BOLD responses were calculated across all trials within a given subject. **a)** Average response for high-intensity stimulation separated by arousal state. In the LGN, we observed consistent positive BOLD responses for high-intensity stimulation that was similar across all arousal states (wake vs. N1: $t(26)$=-0.351, $p$=0.728; wake vs. N2: $t(18)$=-0.119, $p$=0.906; N1 vs. N2: $t(20)$=0.482, $p$=0.635). However, in visual cortical ROIs, we observed a suppression of BOLD responses from wake to N2. From wake to N1 and N2 there was a suppression of BOLD responses in V1 (wake vs. N1: $t(26)$=2.789, $p$=0.009; wake vs. N2: $t(18)$=1.945, $p$=0.067; N1 vs. N2: $t(20)$=0.479, $p$=0.637), V2 (wake vs. N1: $t(26)$=2.433, $p$=0.022; wake vs. N2: $t(18)$=2.281, $p$=0.035; N1 vs. N2: $t(20)$=0.281, $p$=0.781), and V3 (wake vs. N: $t(26)$=2.062, $p$=0.059; wake vs. N2: $t(18)$=1.437, $p$=0.184; N1 vs. N2: $t(20)$=0.243, $p$=0.813). In V1 the difference between wake vs. N2 is marginally significant and in V3 the difference between wake vs. N2 is not statistically significant. **b)** Average response for low-intensity stimulation separated by arousal state. There were no significant differences in responses to low-intensity stimulation between any arousal states (all $p$>0.05). Individual subjects are shown as transparent dots. Error bars represent standard error across participants.

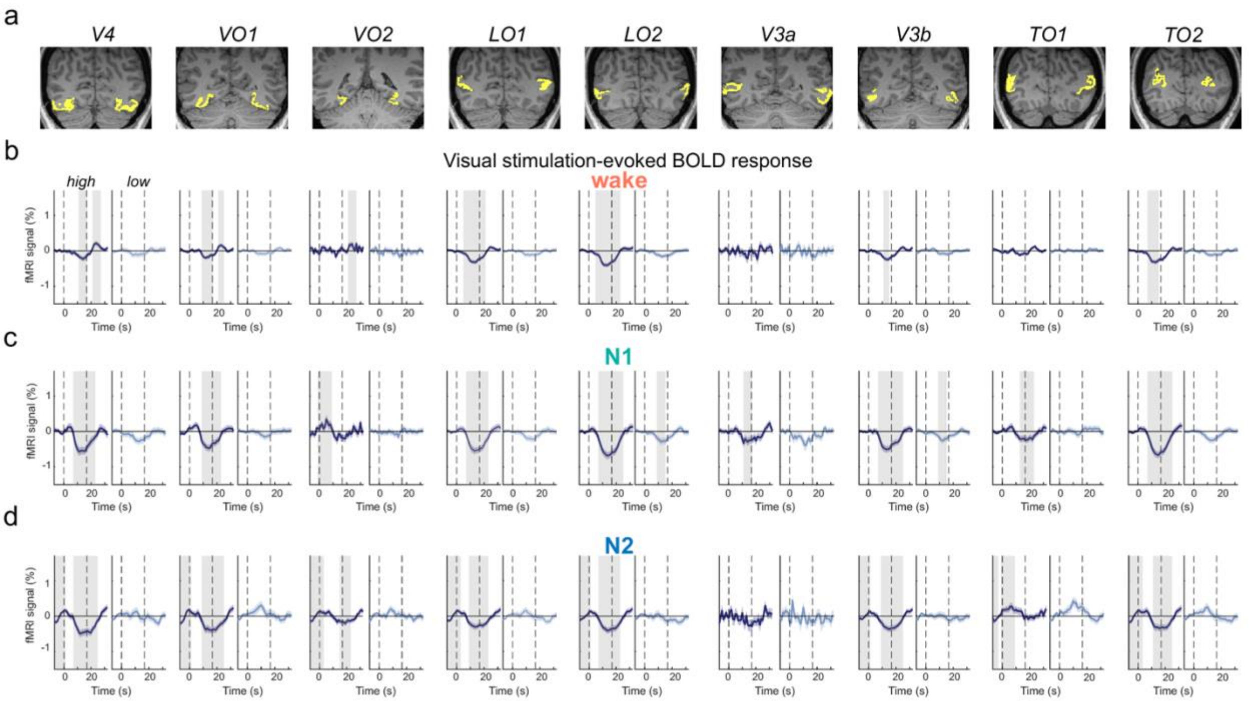


**Fig. S3. Visual stimulation during N1 and N2 sleep produces negative responses throughout most visuocortical regions. a)** Other visuocortical ROIs from the Bayesian segmentation algorithm (*84*) are shown in coronal slices in an example subject. **b-d)** Stimulus-locked BOLD responses from other visuocortical ROIs separated by temporal contrast modulation (high-intensity: dark blue; low-intensity: light blue) and sleep-wake state during stimulus presentation. Significant differences between high- and low-intensity responses with baseline were calculated with sliding window time-binned linear mixed effects models (with random effect of subject) and corrected for number of time bins, number of ROIs, and number of arousal states using FDR correction (*93*). High-intensity stimulation during wake (**b**), N1 (**c**) and N2 (**d**) evoked significant deactivation across most higher-order visuocortical regions (grey bars indicate $p<0.05$; time-binned linear mixed effects model). All higher-order visual cortical regions, except for V3a ($p>0.05$), displayed a significantly reduced BOLD response during N1 and N2 compared to wakefulness ($p<0.05$), indicating enhanced suppression across the visual cortex during sleep. No visual cortical regions showed significant differences in BOLD responses between N1 and N2 seep ($p>0.05$). Shading is standard error. Dashed vertical lines represent stimulus onset (left line) and offset (right line).

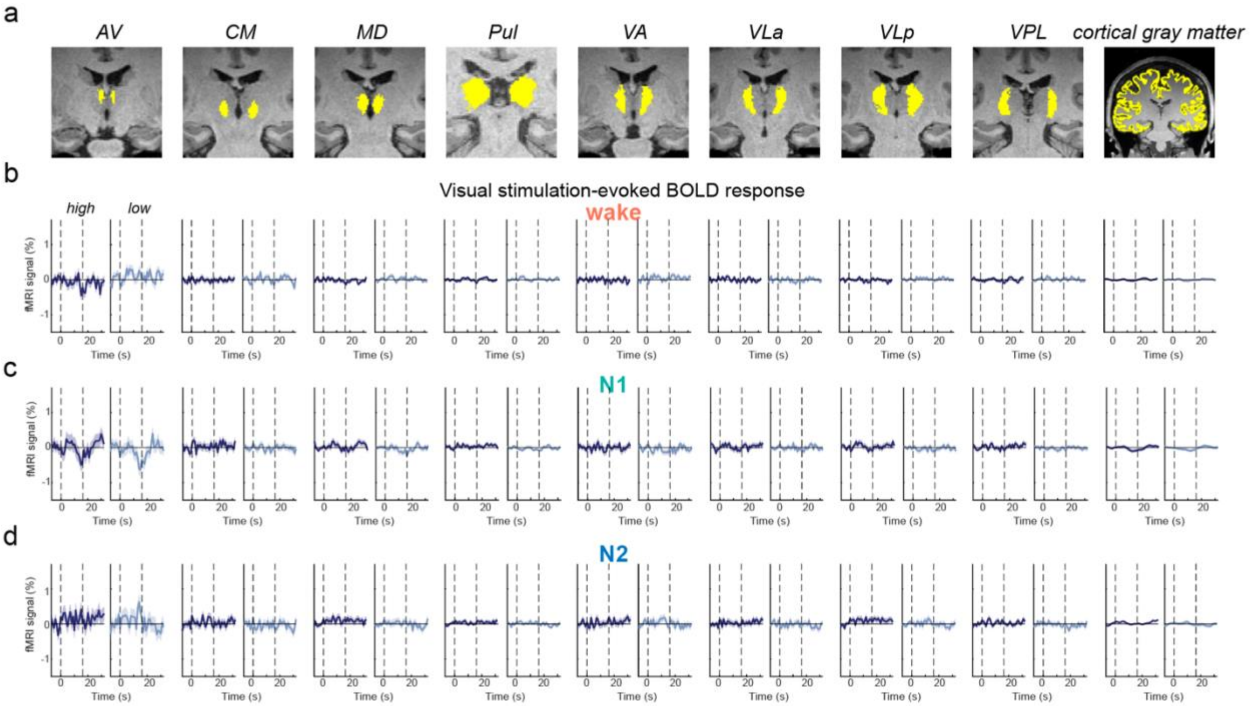


**Fig. S4. Visual stimulation does not evoke significant responses in thalamic nuclei other than the LGN, nor across the whole cortical gray matter across arousal states. a)** Thalamic nuclei and cortical gray matter regions of interest are shown in coronal slices in an example subject. **b-d)** Stimulus-locked BOLD responses were averaged across all cortical ROIs using the FreeSurfer whole-brain segmentation (*83*) and from all thalamic nuclei from FreeSurfer's thalamic segmentation algorithm (*87*), separated by stimulus intensity (high-intensity: dark blue; low-intensity: light blue) and sleep-wake state during stimulus presentation. Significant differences between high- and low-intensity responses with baseline were calculated with sliding window time-binned linear mixed effects model (with random effect of subject) and corrected for number of time bins, number of ROIs, and number of arousal states using FDR correction (*93*). During wakefulness (**b**), N1 (**c**), and N2 (**d**) no significant responses were identified across thalamic nuclei (except for LGN in Fig. 2) nor across the whole cortical gray matter (all $p$'s>0.05). Shading is standard error. Dashed vertical lines represent stimulus onset (left line) and offset (right line).

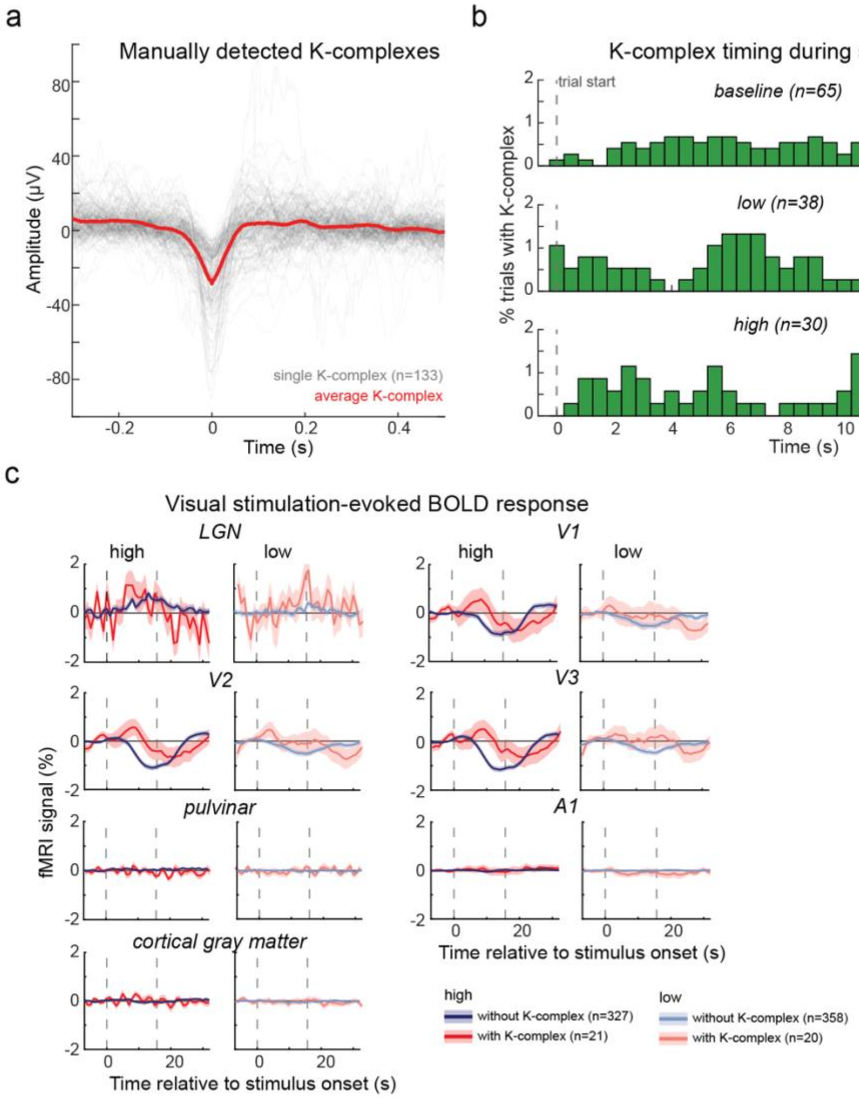


**Fig. S5. K-complexes do not underlie the negative visuocortical BOLD during high-intensity visual stimulation during sleep. a)** K-complexes were manually detected according to AASM criteria (*22*). We detected 133 K-complexes in frontal and central electrodes. Individual K-complexes are shown in light gray and the average K-complex is in red. Some detected K-complexes are small in amplitude relative to those observed from EEG outside of an MRI scanner; however, per AASM definitions (*22*) they were identified by characteristic morphology rather than amplitude, and attenuation is expected in simultaneous EEG–fMRI recordings using average referencing. **b)** Peri-stimulus time histogram (500 ms bins) of K-complex events. The timing of K-complexes relative to trial onset shows no clear relationship between trial onset and K-complex occurrence across all trial types. Dashed vertical lines indicates stimulus duration. **c)** Visual stimulation-evoked BOLD responses for high- and low-intensity trials during N2 sleep were calculated separately for those trials where a K-complex occurred anywhere during the 16 second trial (high-intensity: $n$=21; low-intensity: $n$=20) and for those without a K-complex during the trial (high-intensity: $n$=327; low-intensity: $n$=358). To determine if the presence of a K-complex elicited a significantly different BOLD response during visual stimulation, time-binned linear mixed effects models were computed (5 TR window, 1 TR step size) and p-values were corrected for number of bins and number of ROIs using FDR correction (*93*). There were no significant differences in any ROIs BOLD responses with and without the occurrence of a K-complex (all $p$'s>0.05). Since the vast majority of trials did not elicit a K-complex, K-complexes could not explain the negative fMRI responses. Shading is standard error. Dashed vertical lines represent stimulus onset (left line) and offset (right line).

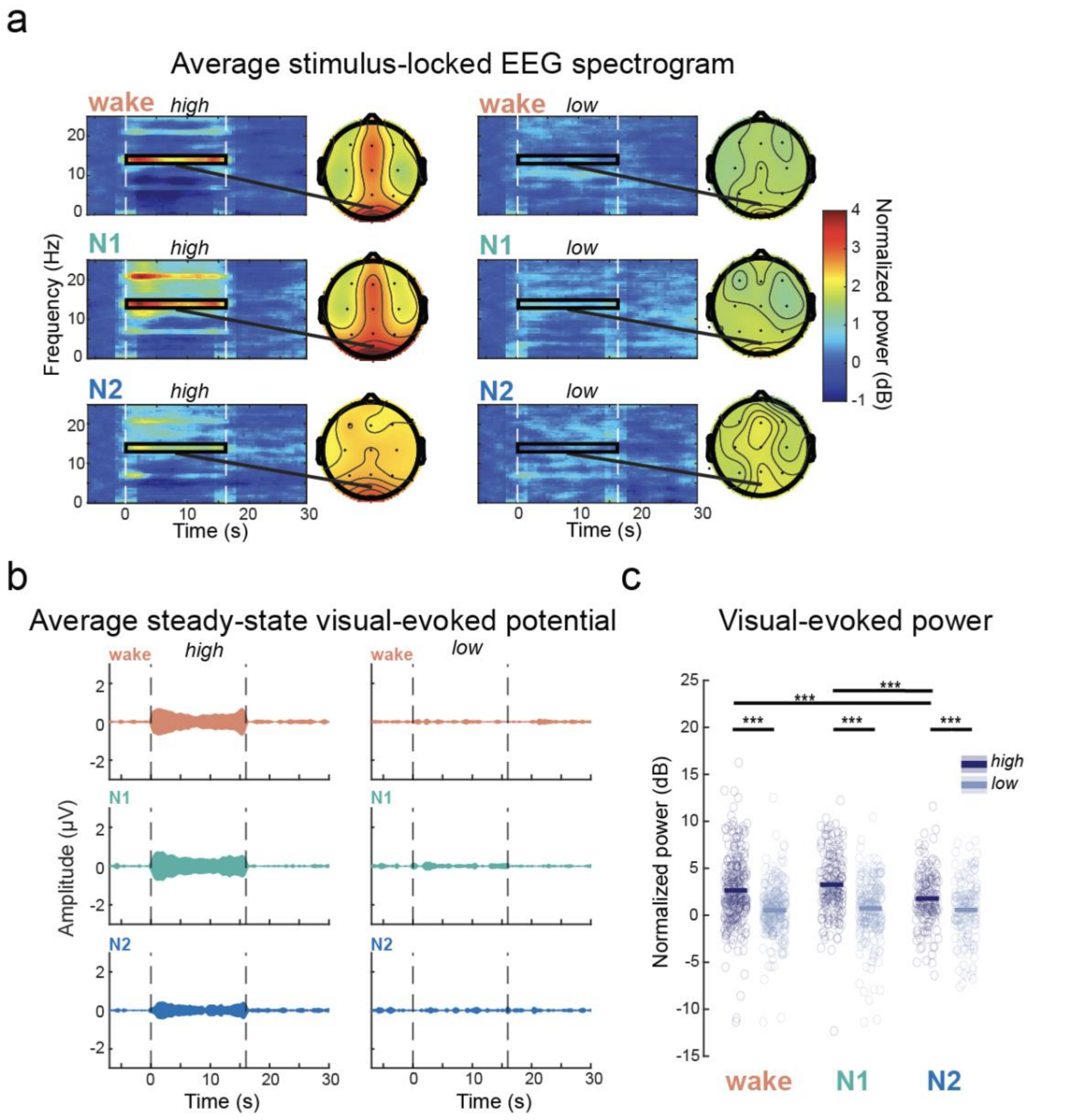


**Fig. S6. Occipital EEG visual-evoked potential is present, but attenuated, during N2 sleep. a)** Average stimulus-locked EEG spectrogram (channel Oz) separated by arousal state and stimulus intensity. The spatial topography of the average visual evoked potential (14 Hz power) across the 16s stimulus presentation for the corresponding arousal state and stimulus condition is displayed across all electrodes to the right of each spectrogram. Stimulus presentation is represented by white dashed lines at 0s and 16s. **b)** Average steady-state visual evoked potential from occipital EEG channels across arousal states and stimulus intensities. For each trial, the EEG data was bandpass filtered (13.5-14.5 Hz) and baseline normalized to the average amplitude 4 seconds prior to stimulus onset. Stimulus duration is represented by grey dashed lines at 0s and 16s. **c)** 14 Hz visual-evoked power averaged across the 16s stimulus period from occipital EEG channels. ANOVA revealed a significant effect of arousal state ($F(2)=5.43$, $p=0.005$), stimulus intensity ($F(1)=82.43$, $p<1.0e^{-12}$), and an interaction between stimulus intensity and arousal state ($F(2)=3.98$, $p=0.016$). Pairwise comparisons revealed significantly greater 14 Hz power for high- compared to low-intensity trials across all arousal states (wake: $t(592)=6.67$, $p<1.0e^{-11}$; N1: $t(369)=6.94$, $p<1.0e^{-11}$; N2: $t(280)=2.86$, $p=0.005$). During N2 sleep, 14 Hz power during high-intensity trials was significantly reduced compared with wake ($t(451)=2.24$, $p=0.025$) and N1 sleep ($t(310)=4.517$, $p<1.0e^{-5}$).

| subject | % *wake* | % *N1* | % *N2* | *# trials wake* | | *# trials N1* | | *# trials N2* | |
|---|---|---|---|---|---|---|---|---|---|
| | | | | high | low | high | low | high | low |
| *1* | 23% | 34% | 43% | 8 | 9 | 18 | 17 | 20 | 21 |
| *2* | 49% | 51% | 0% | 21 | 17 | 12 | 24 | 0 | 0 |
| *3* | 55% | 40% | 5% | 38 | 24 | 18 | 25 | 1 | 1 |
| *4* | 79% | 21% | 0% | 39 | 44 | 11 | 7 | 0 | 0 |
| *5* | 64% | 32% | 4% | 12 | 12 | 2 | 1 | 0 | 0 |
| *6* | 74% | 21% | 5% | 48 | 39 | 9 | 10 | 3 | 1 |
| *7* | 51% | 44% | 5% | 24 | 21 | 13 | 21 | 3 | 1 |
| *8* | 70% | 29% | 1% | 37 | 37 | 17 | 12 | 0 | 1 |
| *9* | 34% | 37% | 29% | 14 | 13 | 13 | 16 | 13 | 9 |
| *10* | 2% | 53% | 45% | 0 | 0 | 27 | 19 | 18 | 25 |
| *11* | 31% | 32% | 37% | 7 | 7 | 6 | 10 | 23 | 15 |
| *12* | 36% | 26% | 38% | 16 | 14 | 4 | 13 | 14 | 16 |
| *13* | 15% | 18% | 67% | 2 | 0 | 6 | 11 | 36 | 36 |
| *14* | 55% | 10% | 35% | 10 | 13 | 1 | 2 | 11 | 9 |
| *15* | 59% | 32% | 9% | 32 | 36 | 10 | 16 | 3 | 2 |

**Table S1. Summary statistics for trials across subjects.** Percentage of resting-state run from wake, N1, and N2 for each subject based on clinical scoring criteria (*22*). No data was observed during NREM stage 3 (N3) or REM sleep. The number of trials across wake, N1, and N2 is also displayed for each trial type per subject.
